# Non-ionotropic signaling through the NMDA receptor GluN2B carboxy terminal domain drives morphological plasticity of dendritic spines and reverses fragile X phenotypes in mouse hippocampus

**DOI:** 10.1101/2024.12.15.628559

**Authors:** Stephanie A. Barnes, Aurore Thomazeau, Peter S.B. Finnie, Maxwell J Heinrich, Arnold J. Heynen, Noburu H. Komiyama, Seth G.N. Grant, Frank S. Menniti, Emily K. Osterweil, Mark F. Bear

## Abstract

It is well known that activation of NMDA receptors can trigger long-term synaptic depression (LTD) and that a morphological correlate of this functional plasticity is spine retraction and elimination. Recent studies have led to the surprising conclusion that NMDA-induced spine shrinkage proceeds independently of ion flux and requires the initiation of *de novo* protein synthesis, highlighting an unappreciated contribution of mRNA translation to non-ionotropic NMDAR signaling. Here we used NMDA-induced spine shrinkage in slices of mouse hippocampus as a readout to investigate this novel modality of synaptic transmission. By using selective pharmacological and genetic tools, we find that structural plasticity is dependent on the ligand binding domain (LBD) of GluN2B-containing NMDA receptors and that metabotropic signaling occurs via the GluN2B carboxyterminal domain (CTD). Disruption of signaling by replacing the GluN2B CTD with the GluN2A CTD leads to increased spine density, dysregulated basal protein synthesis, and epileptiform activity in area CA3 reminiscent of phenotypes observed in the *Fmr1^−/y^* model of fragile X syndrome. By crossing the *Fmr1^−/y^* mice with animals in which the GluN2A CTD has been replaced with the GluN2B CTD, we observe a correction of these core fragile X phenotypes. These findings suggest that non-ionotropic NMDAR signaling through GluN2B may represent a novel therapeutic target for treatment of fragile X and related causes of intellectual disability and autism.

## INTRODUCTION

N-methyl-d-aspartate receptors (NMDARs) play a crucial role in refining neural circuits that are established during development and modified during experience-dependent learning. These excitatory glutamatergic receptors mediate dynamic changes in synaptic connectivity, which include the controlled regulation of dendritic spine formation and enlargement, as well as spine retraction and elimination. Bidirectional changes in dendritic spine structure coincide with changes in synaptic strength. Enlargement of dendritic spines accompanies long-term potentiation (LTP), whereas spine shrinkage is observed after long-term depression (LTD). Considerable evidence suggests that perturbations in the maturation and plasticity of neuronal connections underlie neurological disorders associated with cognitive impairments. In agreement, numerous mutations in NMDAR subunits have been identified in patients with intellectual disability (ID), autism spectrum disorder (ASD), and epilepsy^1–3^. These findings indicate that NMDAR dysfunction may play a prominent role in the pathophysiology of neurodevelopmental disorders.

In the CA1 region of the hippocampus, NMDAR-dependent LTD and spine shrinkage can be induced by brief application of the agonist NMDA^4^, low-frequency stimulation of identified input pathways^5^, or by uncaging glutamate at single spines^6^. From these studies it has emerged that NMDARs can trigger intracellular events independently of ion permeation^7–9^. Although there are conflicting data on the requirement for Ca^2+^ flux to trigger NMDAR-LTD^6,10–15^, numerous groups have consistently shown that structural plasticity is solely dependent on agonist binding, and occurs even in the presence of co-agonist and ion channel blockers^6,16^. Recent studies from this laboratory, in which the functional and structural effects of brief exposure to the receptor agonist NMDA were monitored simultaneously, confirmed that LTD and spine shrinkage mechanistically diverge^16^. NMDAR-LTD was abolished in the presence of the ion channel blocker MK-801, and by 7-CK an antagonist at the co-agonist binding site, while spine shrinkage remained intact. Together these findings indicate that the NMDAR can signal in both an ionotropic and non-ionotropic mode to trigger LTD and spine shrinkage, respectively. Non-ionotropic NMDAR signaling was originally described as “metabotropic”^10,17^, so we have used the term “mNMDAR” to refer to this type of signaling^18^.

Further examination of mNMDAR signaling revealed that spine shrinkage was dependent on the activation of mTORC1 and the synthesis of new proteins^16^. Dysregulated synaptic protein synthesis and altered spine structure are features of fragile X syndrome, a neurodevelopmental disorder characterized by ID and ASD, and caused by loss of the fragile X messenger ribonucleoprotein (FMRP)^19,20^. This structural readout of mNMDAR function was therefore investigated in the *Fmr1^−/y^* mouse model of fragile X. Although the magnitude of spine shrinkage in response to NMDA appeared intact, it proceeded in the presence of cycloheximide, consistent with the conclusion that the local protein synthesis that normally gates spine plasticity is dysregulated in this disorder^16^.

There is much left to learn about mNMDARs and how they might contribute to pathophysiology in fragile X. In this study we have used spine shrinkage in response to NMDA as an assay to dissect the GluN2 subtype and structural motifs that transduce agonist binding to the initiation of mNMDAR signaling. The data show that ligand binding to the GluN2B subunit is obligatory, and that signaling is mediated via the GluN2B carboxyterminal domain (CTD). Eliminating mNMDAR signaling by replacing the GluN2B CTD with the GluN2A CTD phenocopies several core aspects of fragile X pathophysiology in the hippocampus; namely, increased density of dendritic spines, an elevated rate of bulk basal protein synthesis, exaggerated LTD mediated by G-protein coupled metabotropic glutamate receptors (mGluR-LTD) in area CA1, and increased epileptiform activity in area CA3. Conversely, by crossing *Fmr1^−/y^* mice with animals in which mNMDAR signaling is augmented by replacing the GluN2A CTD with the GluN2B CTD, we find correction of protein synthesis, mGluR-LTD, and CA3 epileptiform activity. Together, the results suggest that GluN2B might be a novel therapeutic target in fragile X.

## METHODS

### Animals

Thy1-GFP mice were obtained from Jackson Laboratories, Maine, USA (stock # 003025 and # 011070, respectively) and maintained on a congenic C57BL/6J background. Homozygous floxed GluN2A (Grin2A^fl/fl^) and GluN2B (Grin2B^fl/fl^) were obtained from the Nicoll laboratory at UCSF^21,22^. The GluN2A KO line was acquired from the Constantine-Paton laboratory at MIT and transgenic knockin lines GluN2A^(2BCTD)^ and GluN2B^(2ACTD)^ mice were obtained from Komiyama/Grant laboratories at the University of Edinburgh^23^ and crossed with Thy1-GFP line. Generation of double mutant lines involved crossing *Fmr1^+/−^* females with *GluN2A^2B(CTR)^ ^+/−^* males with only first-generation litters taken. Experimental cohorts consisted of female and male littermates, except *Fmr1* KO lines, that were P25-35 at the time of experiments. Mice were group housed with littermates and maintained on a 12:12h light:dark cycle. All experiments were performed blind to genotype using age-matched littermate controls during the light phase. The Institutional Animal Care and Use Committee at Massachusetts Institute of Technology approved all experimental techniques.

### CA1 viral infusions

Juvenile mice (P25-30) were anaesthetized and received subcutaneous injections of preoperative analgesics (buprenorphine (0.1 mg/kg, meloxicam; 1 mg/kg, and 1% lidocaine) before being head-fixed on a stereotaxic frame. Craniotomies were performed over CA1 where 20 short pulses of 20 nl of herpes simplex virus (HSV) expressing a cre-eGFP fusion protein (HSV-eGFP-cre) or HSV-eGFP as a control (∼8.75×10^8^ particles/ml) was injected at a rate of 43 nl/minute at 2-3 sites using a Nanoject III (Drummond Scientific) and a beveled glass injection pipette. After surgery mice received post-operative analgesics and were returned to their home cage. Around 5-7 days, mice were sacrificed for recordings. Cre expression was generally limited to a sparse population of CA1 pyramidal cells in the dorsal hippocampus.

### Preparation of Hippocampal Slices

At MIT, mice were anesthetized through isoflurane inhalation (AErrane; Baxter Pharmaceuticals), sacrificed and hippocampi were isolated. Acute dorsal hippocampal slices (350 μm thick) were prepared using a vibratome (Leica Microsystems) in ice-cold dissection buffer containing (in mM): NaCl 87, sucrose 75, KCl 2.5, NaH_2_PO_4_ 1.25, NaHCO_3_ 25, CaCl_2_ 0.5, MgCl_2_ 7, ascorbic acid 1.3, and D-glucose 10 (saturated with 95% O_2_,5% CO_2_). For LTD and spine shrinkage experiments, the CA3 region was removed. Slices were recovered in artificial cerebrospinal fluid (ACSF) containing (in mM): NaCl 124, NaH_2_PO_4_ 1.2, KCl 5, NaHCO_3_ 26, glucose 10 CaCl_2_ 2, MgCl_2_ 1 and (saturated with 95% O_2_/5% CO_2_) at 32.5 °C for 30 mins and then recovered at room temperature for 2.5 hrs before recording. At the University of Edinburgh, horizontal hippocampal slices were prepared in ice-cold ACSF containing NaCl, 86; NaH_2_PO_4_, 1.2; KCl, 25; NaHCO_3_, 25; glucose, 20; sucrose, 75; CaCl_2_, 0.5; MgCl_2_, 7; saturated with 95% O_2_, 5% CO_2_. Slices were hemisected and recovered for at least 2 hrs at 32 °C in ACSF containing (in mM): NaCl, 124; NaH_2_PO_4_, 1.2; KCl, 2.5; NaHCO_3_, 25; glucose, 20; CaCl_2_, 2; MgCl_2_; 1, saturated with 95% O_2_, 5% CO_2_.

### Electrophysiology

At MIT, field potential recordings were performed in a submersion chamber, perfused with ACSF (3– 4 ml/min) at 30 °C. fEPSPs were recorded in CA1 stratum radiatum with extracellular electrodes filled with ACSF. Baseline responses were evoked by stimulation of the Schaffer collaterals at 0.033 Hz with a two-contact cluster electrode (FHC, Bowdoin, ME) using a 0.2 ms stimulus yielding 40–60% of the maximal response. Field recordings were filtered at 2 kHz, digitized at 50 kHz and analyzed using pClamp10 (Axon Instruments). The initial slope of the response was used to assess changes in synaptic strength. Functional LTD was quantified by comparing the average response 50– 60 mins after NMDA, to the average of the last 10 mins of baseline. Isolated NMDA-mediated fEPSPs were obtained in ACSF containing Mg^2+^ (0.3 mM) glycine (1 µM), picrotoxin (100 µM) and NBQX (20 µM) before perfusing GluN2 subtype specific blockers conantokin-G (2 µM) and MPX-004 (30 µM).

For extracellular recordings in CA3, an ACSF filled electrode was placed in the stratum pyramidal layer of CA3. Responses were recorded using PClamp10 software in voltage-clamp, amplified x1000, filtered between 300 Hz and 10 kHz and digitized at 25 kHz. A 10-minute baseline was recorded in ACSF before perfusing in either the GABA_A_ antagonist bicuculline (50 µM) or the group 1 mGluR_1/5_ agonist DHPG (50 µM). Voltage traces were Butterworth filtered between 300 and 1000 Hz before being downsampled by a factor of 10. The spike threshold was determined by calculating the median absolute deviation and multiplying by a factor of 5. Following spike detection, spikes were summed across 100 msec bins to reduce computational expense. To detect bursting events, a 500 msec sliding window was pulled across the entirety of the recording, from 5 to 60 minutes following the baseline period. A burst was identified if the spike rate within the window exceeded 8 Hz and was terminated in subsequent windows when the spike rate fell below 4 Hz.

At the University of Edinburgh, extracellular recordings were performed by placing slices in a submersion chamber heated to 30 ^◦^C (Fine Science Tools) and perfused with pre-oxygenated ACSF at a rate of 4 ml/min. mGluR-dependent LTD was induced in CA1 by perfusing the slice with the group 1 mGluR agonist DHPG (50 µM, 5 mins). Waveform data was collected using WinLTP 1999-2009 (WinLTP Ltd., University of Bristol), amplified 1000 times (npi electronics), filtered at 1.3 kHz and digitized (National Instruments) at 20 kHz. The data was exported to Microsoft Excel and the magnitude of LTD was calculated from the last 10 minutes of the recording relative to the pre-drug baseline.

### Two-photon laser-scanning microscopy

Time-lapse fluorescence two-photon imaging was performed using a Prairie Technologies Ultima system attached to an Olympus BX-51WI that was equipped with a mode-lock femtosecond-pulse Ti:Sapphire laser (Chameleon, Coherent). Experiments have been previously described in detail in^16^. Briefly, GFP was excited at 930 nm and images were taken with a 60X 0.9 NA objective lens, and a digital zoom of x 5.60 every 4 mins.

### Metabolic Labeling

Hippocampi were rapidly dissected from rodents and transverse hippocampal slices were prepared using a Stoelting tissue slicer in ice-cold artificial cerebrospinal fluid (ACSF) containing: (in mM): NaCl, 124; NaH_2_PO_4_, 1.25; KCl, 3; NaHCO_3_, 26; glucose, 10; CaCl_2_, 2; MgCl_2_, 1, saturated in 95% O_2_ and 5% CO_2_. Hippocampal slices were left to recover for at least 4 hours at 32°C in oxygenated ACSF. Transcription was blocked with actinomycin D (ActD, 25 µM; Tocris) ± drug for 30 minutes then slices were metabolically labelled in ^35^S-Methionine/Cysteine express protein labeling mix (Perkin Elmer) ± drug for 30 minutes. Samples were homogenized in buffer containing: 10 mM HEPES pH 7.4, 2 mM EDTA, 2 mM EGTA, 1% triton X-100 (Sigma-Aldrich), protease and phosphatase inhibitors. Proteins were precipitated with TCA (Sigma) and resolubilised. Counts per minute (CPM) were quantified and normalised to overall protein using a DC protein assay kit II (Biorad).

### Pharmacological agents

NMDAR-dependent LTD and spine shrinkage was induced by the acute application of NMDA (20 µM, 3 min). For pharmacological experiments, slices were pre-incubated in the respective vehicle/drug for 40 mins prior to recording/imaging, and then continually perfused throughout the entire experiment. NMDA, MK-801, picrotoxin, glycine, NBQX and DHPG (U.of.E) were purchased from Sigma. Conantokin-G, DHPG (MIT), and bicuculline, were purchased from Tocris Biosciences, while MPX-004 (30 µM) was purchased from Alomone.

### Image analysis

Throughout the baseline and LTD recording, a 512×512 pixel XY-scanned Z-series was taken every 4 minutes, at 1 µm of tissue depth for 15 µm of depth in total. Maximal fluorescence intensity of the Z-Series was summed to obtain a Z-stack for every time point and collated to form a movie montage, aligned using the StackReg function in Fiji/ImageJ (by Rasband, W.S., U. S. National Institutes of Health). Typically, 15 spines on multiple dendritic regions were tracked throughout the recording by outlining a 20 x 20 pixel region of interest (ROI). Within this ROI, the total integrated fluorescence intensity of the green was calculated using ImageJ. Background fluorescence was tracked at 3 locations and subtracted, then overall fluorescence fluctuations was corrected by measuring the intensity over time at 4 locations. These values were then taken to be proportional to spine volume. Only experiments showing <7% of drift variation during the baseline period, calculated by fitting a linear regression line for the 30 min of baseline were utilized.

### Statistical analyses

All data is expressed as a mean ± S.E.M with statistical significance calculated using Prism 9.0 (GraphPad software) with a confidence level set at 95%. For single comparisons between genotype or treatment, significance was calculated by Student’s *t*-test or one-way ANOVA with Bonferroni *post-hoc* test. Multiple comparisons (two genotypes/two treatments/genotype x treatment) were analyzed using two-way ANOVA with *post-hoc* Student’s *t*-test. Pharmacological or genetic experiments were statistically compared to their corresponding vehicle or WT controls. Experiments were performed blind to genotype and statistics were performed using N as animal.

## RESULTS

In hippocampal pyramidal neurons, NMDARs are tetramers consisting of two obligatory GluN1 and two GluN2 (A and B) subunits that co-assemble in to a di- or tri-heteromeric arrangement. Although GluN2B predominates at birth and is expressed throughout adulthood, GluN2A levels gradually increase during postnatal development and impose a distinct and dominant kinetic profile on the NMDAR assembly^24,25^. Since GluN2A/2B subunits have varied distributions and physiological roles, an important objective is to determine the subtype specificity of mNMDAR signaling and understand how it elicits structural plasticity at CA1 synapses. Study of the respective contribution of the GluN2 subtype is complex because deletion of *Grin2B* is lethal from birth and widespread loss leads to disruption in network activity^26,27^. Therefore, in addition to utilizing a constitutive *Grin2A* KO mouse line, we employed genetically engineered mice carrying conditional knockout alleles for GluN2A (*Grin2A^fl/fl^*) and GluN2B (*Grin2B^fl/fl^*) to determine the subtype specificity of mNMDAR signaling.

We simultaneously examined the contribution of the GluN2A subtype to NMDAR-dependent LTD and spine shrinkage in transverse hippocampal slices prepared from the progeny of *Grin2A* KO mice crossed with a Thy1-GFP line (P25-35). These mice express GFP in a sparse subset of neurons and using two-photon time-lapse fluorescence imaging we were able to visualize dendritic spines along isolated apical dendrites of CA1 pyramidal neurons in stratum radiatum. In addition, we performed extracellular recordings by stimulating the Schaffer collateral pathway every 30 seconds and recorded the corresponding field excitatory postsynaptic potentials (fEPSPs) from the region surrounding the dendrite of interest. To induce NMDAR-dependent LTD and spine shrinkage, we briefly applied NMDA (20 μM, 3 min), which is known to trigger a saturating and persistent reduction in the fEPSP slope and spine volume^4,18^.

### GluN2B di-heteromers trigger non-ionotropic NMDAR signaling and the retraction of dendritic spines

We observed that the acute application of NMDA induced a persistent depression of fEPSP responses in WT Thy1-GFP slices that was significantly reduced, but not fully abolished, in *Grin2A* KO Thy1-GFP mice (**Fig. 1A_i_**). In contrast, the NMDA-triggered reduction in spine volume was indistinguishable between *Grin2A* KO and WT Thy1-GFP slices (**Fig. 1A_ii_**). These findings suggest that GluN2A-containing assemblies play a role in functional NMDAR-LTD but are not required for the induction of spine shrinkage. Therefore, the GluN2B di-heteromers may be sufficient to initiate the non-ionotropic signaling that underlies NMDA-induced structural plasticity at CA1 synapses.

**Figure 1:**
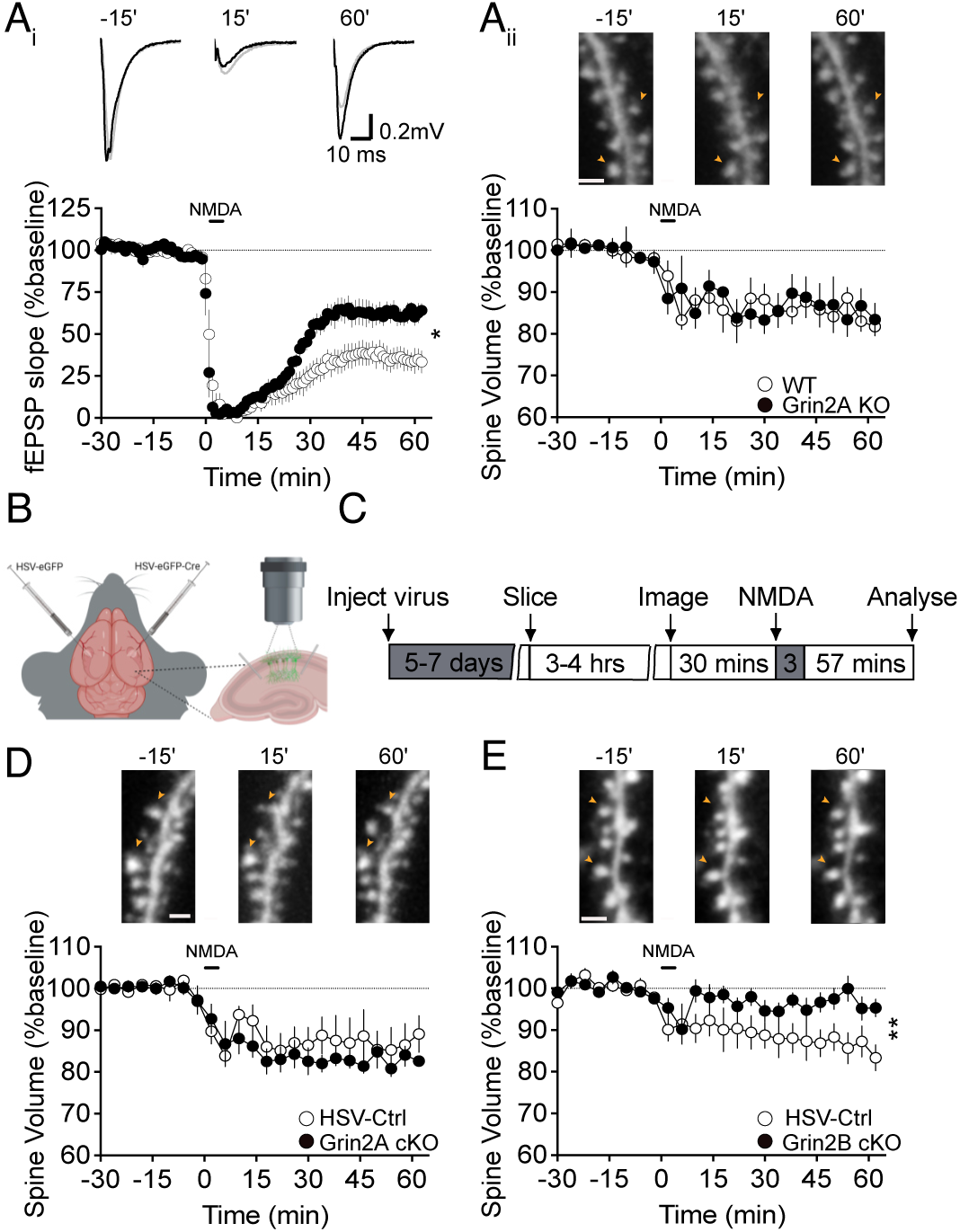
Spine structural plasticity triggered by non-ionotropic NMDAR signaling depends on the GluN2B subunit. (**A_i-ii_**) Extracellular field recordings and time-lapse two-photon imaging were simultaneously performed in CA1 stratum radiatum of acute hippocampal slices prepared from *Grin2A* KO Thy1-GFP mice and WT littermates. Representative fEPSP traces and images are shown 15 mins before, 15 mins and 60 mins after LTD induction. (**A_i_**) Brief bath application of NMDA (20 μM; 3 mins) induces a persistent reduction in the slope of the fEPSP in WT Thy1-GFP slices that is significantly impaired in *Grin2A* KO Thy1-GFP slices (WT: 35 ± 6%, *n* = 8; *Grin2A* KO: 62 ± 5%, *n* = 5; *p* = 0.01, unpaired *t*-test). (**A_ii_**) In contrast, NMDA triggered a decrease in spine volume in WT Thy1-GFP slices persisted in the absence of the *Grin2A* allele (WT: 84 ± 2%, *n* = 8; *Grin2A* KO: 85 ± 5%, *n* = 5; *p* = - 0.87, unpaired *t*-test) indicating that the GluN1/GluN2B di-heteromeric receptors are sufficient to support mNMDAR signaling. (**B**) Schematic of HSV injections into the dorsal hippocampus of floxed *Grin2A* and *Grin2B* mice. (**C**) Experimental timeline of conditional knockout (cKO) of *Grin2A* and *Grin2B* in the dorsal hippocampus of floxed *Grin2A* and *Grin2B* mice through injections of either HSV-eGFP control or HSV-eGFP-Cre. (**D**) Conditional deletion of the *Grin2A* gene has no effect on structural plasticity in response to NMDA receptor activation (HSV Control: 87 ± 5%, *n* = 4; HSV-Cre-*Grin2A* cKO 83 ± 1%, *n* = 7; *p* = 0.36, unpaired *t*-test). (**E**) Loss of *Grin2B* significantly impairs NMDA-induced spine shrinkage (HSV Control: 86 ± 3%, *n* = 6; HSV-Cre-*Grin2B* cKO: 97 ± 2%, *n* = 10; *p* = 0.0097, unpaired *t-*test).

To test this hypothesis, juvenile *Grin2A* and *Grin2B* floxed mice (P25) received a transcranial injection into the dorsal hippocampus of a HSV expressing cre-eGFP fusion protein or HSV-eGFP control (**Fig. 1B**). After 5-7 days, hippocampal slices were prepared and imaged (**Fig. 1C**). Consistent with previous reports, we observed sparse eGFP signal throughout CA1 indicating that the HSV-cre-eGFP eliminated the floxed *Grin2A or 2B* allele in a small population of pyramidal neurons in CA1^21^. In agreement with our findings from *Grin2A* KO Thy1-GFP mice, the acute application of NMDA led to a comparable reduction in spine volume in conditional *Grin2A^fl/fl^* and control mice (**Fig. 1D**). However, conditional deletion of *Grin2B* postnatally, which spares only the GluN2A di-heteromeric receptors, abolished spine shrinkage at CA1 synapses (**Fig. 1E**). Thus, the GluN2B-containing receptors are obligatory and sufficient to induce mNMDAR signaling. Together, these findings indicate that GluN2A- and GluN2B-containing di- and tri-heteromeric assemblies contribute differentially to NMDAR-dependent functional and structural plasticity at CA1 synapses.

### Pharmacology of NMDAR-mediated functional and structural plasticity indicates that agonist binding to GluN2B is critical for non-ionotropic signaling

NMDAR subunits have a similar topology consisting of three transmembrane domains (TMD), one reenterant loop that comprises the channel pore, an extracellular amino-terminal domain (NTD), a ligand binding domain (LBD), and an intracellular carboxyl-terminal domain (CTD) that couples the receptor to intracellular signaling messengers^28^. The GluN2A and GluN2B subtypes show a high degree of similarity in their amino acid structures, particularly in their NTD and TMD (69%), making it difficult to pharmacologically isolate these subunits^29^. However, there are several compounds that show greater selectively towards GluN2A- or GluN2B-containing receptors (**Fig. 2A**).

**Figure 2:**
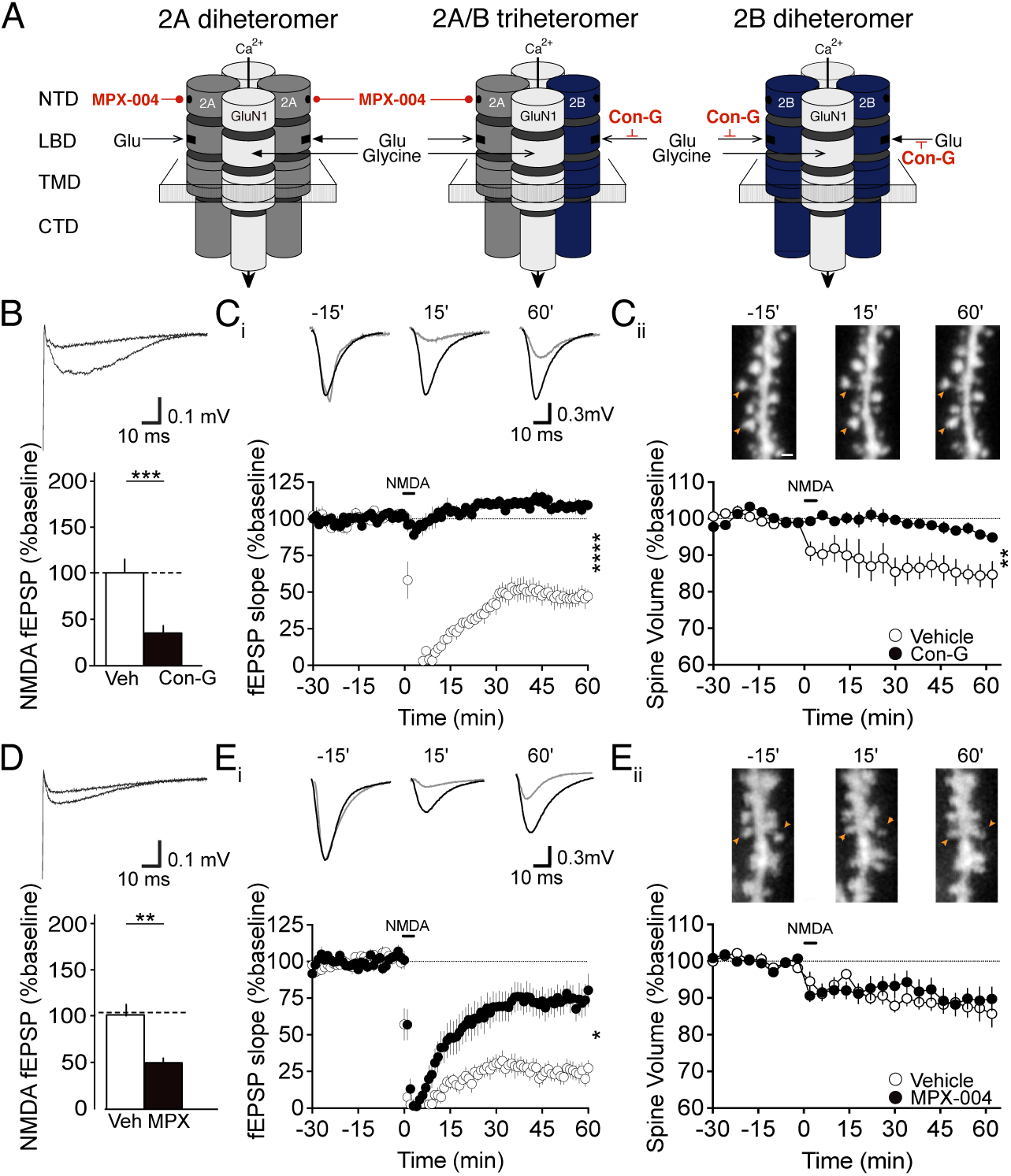
Pharmacology of NMDAR-mediated functional and structural plasticity indicates that agonist binding to GluN2B is critical. (**A**) Schematic of the predominant NMDAR assemblies in hippocampal CA1 and binding sites for GluN2A and GluN2B subtype-selective compounds. The GluN2B competitive antagonist Conantokin-G (Con-G, 2 μM), blocks the glutamate binding site on 2A/2B tri heteromers and 2B di-heteromers. The negative allosteric modulator (NAM) of GluN2A, MPX-004 (40 μM), binds to the N-terminal domain (NTD) of 2A di-heteromers and 2A/2B tri-heteromers. (**B**) Con-G (2 μM) eliminated 66% of the isolated NMDAR-mediated fEPSP (WT vehicle: 100 ± 14%; WT Con-G: 34 ± 6%, *n* = 15; *p* = 0.00016, unpaired *t*-test), abolished both NMDA-induced LTD (**C_i_**; WT vehicle: 46 ± 6%, *n* = 8; WT Con-G: 108 ± 3%, *n* = 10; *p* < 0.0001, unpaired *t*-test) and spine shrinkage at CA1 synapses (**C_ii_**; WT vehicle: 85 ± 3%, *n* = 8; WT Con-G: 96 ± 1%, *n* = 9; *p* = 0.002, unpaired *t*-test). (**D**) MPX-004 (30 μM) caused a 53% block of the isolated NMDAR-fEPSP (WT vehicle: 100 ± 13%; WT MPX-004: 48 ± 4%, *n* = 14; *p* = 0.0013, unpaired *t*-test) and a comparable inhibition of LTD (**Ei**; WT vehicle: 46 ± 7%, *n* = 11; WT MPX-004: 75 ± 11%, *n* = 13; *p* = 0.03; unpaired *t*-test), but had no effect on spine shrinkage (**Eii**; WT vehicle: 87 ± 3%, *n* = 11; WT MPX-004: 89 ± 3%, *n* = 13; *p* = 0.64, unpaired *t*-test). Representative fEPSP traces and images are shown 15 mins before, 15 mins and 60 mins after LTD induction.

Here, we utilized conantokin-G, a competitive antagonist of the GluN2B LBD, and MPX-004, a negative allosteric modulator (NAM) of GluN2A containing NMDARs^30,31^. We observed that blocking the GluN2B LBD with conantokin-G (2 µM), which leaves the LBD of GluN2A di-heteromers intact, eliminated 66% of the isolated NMDAR-mediated fEPSP (**Fig. 2B**) and abolished both NMDA-induced LTD and spine shrinkage at CA1 synapses (**Fig. 2C_i-ii_**). These findings are comparable to those using the non-selective, competitive antagonist D-APV^16^. Therefore, it appears that glutamate must bind to GluN2B-containing di- or tri-heteromeric assemblies to elicit both ionotropic and non-ionotropic NMDAR signaling. The activation of GluN2A di-heteromers alone are not sufficient to drive either NMDA-dependent functional or structural plasticity at CA1 synapses.

Next, we pharmacologically targeted GluN2A using the NAM MPX-004 (30 µM), which strongly inhibits ion flux through the GluN2A di- and tri-heteromeric assemblies^30,32^. We observed that MPX-004 caused a 47% block of the isolated NMDAR-fEPSP (**Fig. 2D**) and a comparable inhibition of LTD (**Fig. 2E_i_**),but had no effect on spine shrinkage (**Fig. 2E_ii_**). Together, the data suggest that GluN2A-containing receptors substantially contribute to the induction of functional LTD, but that spine shrinkage is solely dependent on agonist binding to the LBD of GluN2B-containing receptors.

### NMDARs induce spine shrinkage through the GluN2B carboxyterminal domain

As ligand binding alone can trigger mNMDAR signaling in the absence of ion flux, we hypothesized that it may be transduced through the CTD of GluN2B. While the NTD and TMD domains of GluN2A and GluN2B are highly conserved, their large intracellular CTDs exhibit considerable sequence heterogeneity (∼29% similarity)^29^. It has been previously reported that the GluN2B CTD is crucial for coupling the NMDAR assembly to a large complex of proteins and has distinct effects on synaptic plasticity and behavior relative to the GluN2A CTD^33^. Thus, we speculated that the CTDs may also be the locus for the difference in the abilities of GluN2B and GluN2A NMDAR assemblies to elicit mNMDAR signaling and spine retraction. To test this idea, we utilized a transgenic knockin line, where the exon encoding the CTD of GluN2A is deleted and replaced with GluN2B (GluN2A^2BCTD^, “A2B”; **Fig. 3A**) and vice versa (GluN2B^2ACTD^, “B2A”; **Fig. 3D**). Importantly, the NTD, LBD and TMD are left intact in GluN2A^2BCTD^ and GluN2B^2ACTD^ mice and switching the CTD has been reported to have no functional effect on NMDAR EPSCs or NMDA/AMPAR ratios in the hippocampus^33^. Crossing the GluN2A^2BCTD^ and GluN2B^2ACTD^ mice with the Thy1-GFP line enabled us to then examine structural plasticity downstream of NMDAR activation. We observed that deleting the CTD of GluN2A and replacing it with GluN2B had no impact on spine shrinkage at CA1 synapses (**Fig. 3B**). In contrast, eliminating the CTD of GluN2B and replacing it with GluN2A abolished spine shrinkage in GluN2B^(2ACTD)^ mice relative to WT littermates (**Fig. 3E**). These findings suggest that NMDA triggers mNMDAR signaling through the GluN2B CTD of di-heteromeric assemblies.

**Figure 3:**
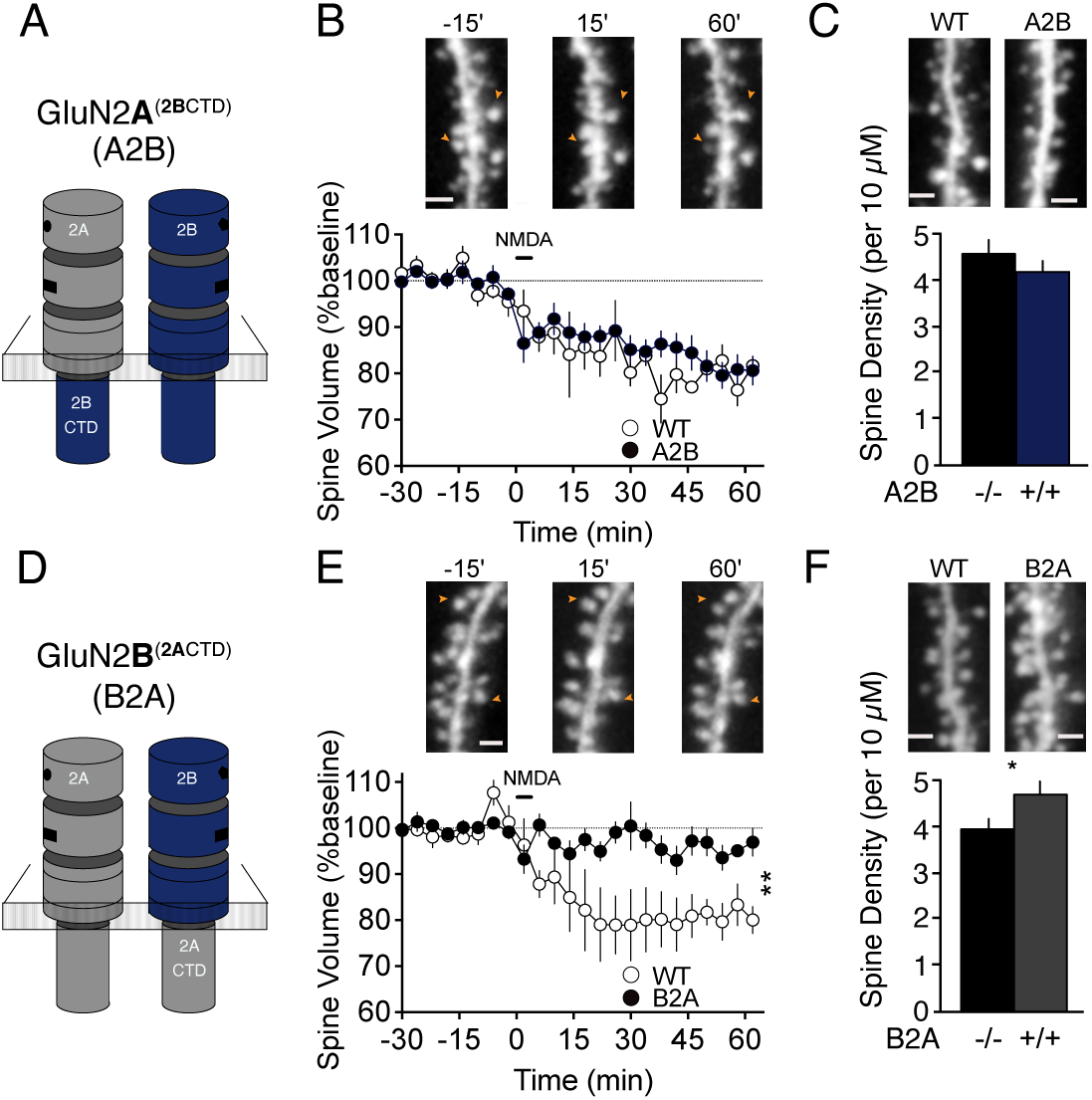
NMDARs induce spine shrinkage through the GluN2B carboxyl terminal domain. (**A**, **D**) Schematic of genetically engineered GluN2A^2BCTD^ (termed A2B) and GluN2B^2ACTD^ (termed B2A) knock-in mouse lines where the exon encoding the CTD of GluN2A is deleted and replaced with GluN2B (GluN2A^2BCTD^) and vice versa (GluN2B^2ACTD^). Representative images are shown 15 mins before, 15 mins and 60 mins after LTD induction (**B**, **C**) Introduction of the GluN2B CTD on GluN2A-containing receptors has no significant effect on structural plasticity (WT: 80 ± 2%, *n* = 4; GluN2A^2BCTD^ HOM: 81 ± 3%, n = 6; *p* = 0.96, unpaired *t*-test) or spine density along apical dendrites of CA1 pyramidal neurones (WT: 4.60 ± 0.64, *n* = 5; GluN2A^2BCTD^ HOM: 4.20 ± 0.21, *n* = 6; *p* = 0.25, unpaired *t*-test). (**E, F**) Switching the GluN2B CTD with 2A leads to a significant impairment in NMDA-induced structural plasticity (WT: 81 ± 3%, *n* = 4, GluN2B^2ACTD^ HOM: 96 ± 2%, *n* = 6, *p* = 0.005, unpaired *t*-test) and a modest increase in spine density in B2A homozygous mice relative to WT littermates (WT: 3.96 ± 0.20, *n* = 5, GluN2B^2ACTD^ HOM: 4.72 ± 0.20, *n* = 7; *p* = 0.029, unpaired *t*-test).

During development, a failure to refine neural circuitry in an activity-dependent manner has been shown to lead to an overabundance of dendritic spines that yields excessive excitatory synapses^34^. Therefore, we speculated whether there may be an abnormality in dendritic spine density in GluN2B^2ACTD^ mice due to the loss of NMDAR-induced spine shrinkage and elimination. To test this hypothesis, we quantified spine density along the apical dendrites of CA1 pyramidal neurons in GluN2A^2BCTD^ and GluN2B^2ACTD^ mice. We observed no significant difference in spine density between GluN2A^2BCTD^ and WT slices (**Fig. 3C**), but in the absence of the 2B CTD there was a subtle, but significant, increase in spine density in GluN2B^2ACTD^ mice relative to WT controls (**Fig. 3F**).

### GluN2B CTD bidirectionally regulates protein synthesis and mGluR-LTD

Previously, we have shown that spine shrinkage is sensitive to the mRNA translational inhibitor cycloheximide and rapamycin, suggesting that mNMDAR signaling may trigger *de novo* protein synthesis via the mTORC1 pathway to support the persistent reduction in spine volume^16^. To directly measure mRNA translation, we performed a metabolic labeling assay in acute hippocampal slices that mimics conditions used to record long-term changes in functional and structural plasticity. Hippocampal slices were recovered in oxygenated ACSF for 4 hrs to allow metabolic processes to recover. Transcription was then blocked with actinomycin D for 30 mins to isolate mRNA translation, before new proteins were labelled with ^35^S-methionine/cysteine protein labeling mix (**Fig. 4A**). To determine whether the activation of NMDARs can drive changes in bulk protein synthesis, we stimulated slices with NMDA (20 µM, 3 mins) or vehicle during protein labeling and then transferred slices to ACSF containing ^35^S methionine/cysteine for a further 27 minutes. We observed that NMDA caused a significant reduction in ^35^S incorporation (**Fig. 4B**), whereas incubating hippocampal slices with the competitive antagonist D-APV, 30 mins prior and throughout protein labeling, led to a significant increase in mRNA translation (**Fig. 4C**). These data demonstrate that targeting the LBD of NMDARs can bidirectionally regulate global protein synthesis rates in the hippocampus.

**Figure 4:**
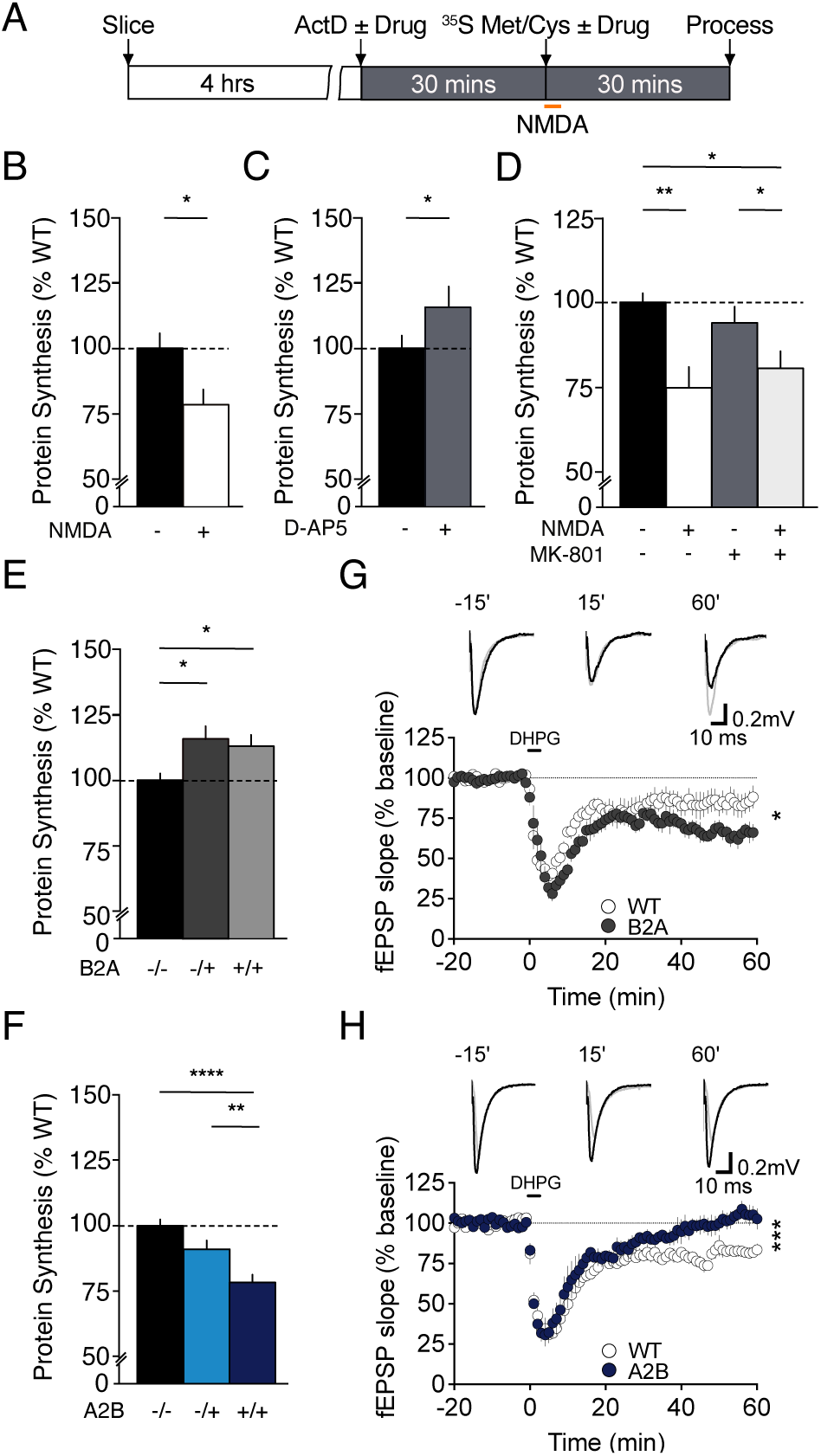
Non-ionotropic NMDAR signaling through the GluN2B CTD negatively regulates bulk protein synthesis and mGluR-LTD. (**A**) Experimental timeline for measuring protein synthesis in acute hippocampal slices. Transverse hippocampal slices were recovered for 4 hrs in ACSF before transcription was blocked with actinomycin D ± drug and newly synthesized proteins radiolabeled with ^35^S methionine/cysteine ± drug. (**B**) The acute application of NMDA (20 μM, 3 mins) leads to a significant decrease in the ^35^S incorporation relative to vehicle-treated slices (vehicle: 100 ± 5%; NMDA: 79 ± 6%; *n* = 8, *p* = 0.014, paired *t*-test). (**C**) In contrast, the non-selective, competitive antagonist D-APV causes a significant increase in protein synthesis in hippocampal slices (vehicle: 100 ± 4%; D-APV: 117 ± 6%; *n* = 12, *p* = 0.046, paired *t*-test). (**D**) The acute application of NMDA leads to a reduction in ^35^S incorporation relative to vehicle treated slices that persisted in the presence of the NMDAR ion channel blocker MK-801 (vehicle: 100 ± 2%; NMDA: 75 ± 6%; MK-801: 97 ± 5%; NMDA/MK-801: 82 ± 5%; *n* = 7, *p* = 0.0018, ANOVA). (**E**) Metabolic labeling reveals bulk protein synthesis is elevated in GluN2B^2ACTD^ (B2A) mice compared to WT littermates (WT: 100 ± 2%, *n* = 13; B2A HET: 116 ± 5%, *n* = 7; B2A HOM: 113 ± 4%, *n* = 8; *p* = 0.0051, ANOVA). (**F**) Metabolic labeling of hippocampal slices reveals the genetic introduction of GluN2B CTD into the GluN2A locus (GluN2A^2BCTD^; A2B) reduces protein synthesis in a gene dose-dependent manner (WT: 100 ± 2%, *n* = 13; A2B HET: 91 ± 3% *n* = 15; A2B HOM: 79 ± 3%, *n* = 15; *p* < 0.0001, ANOVA). (**G**) The acute of application of the group 1 mGluR_1/5_ agonist DHPG (50 μM, 5 mins) induces a long-lasting depression of fEPSPs in CA1 of the hippocampus. Representative fEPSP traces are shown at 15 mins before and 60 mins after LTD induction. Introduction of the GluN2A CTD on GluN2B containing receptors significantly enhanced mGluR-LTD in B2A mice relative to WT littermates (WT: 83 ± 5 %, *n* = 7; B2A HOM: 65 ± 4%, *n* = 6; *p* = 0.017, unpaired *t*-test). (**H**) In contrast, mGluR-LTD is inhibited in the hippocampus of A2B mice relative to WT littermates (WT: 80 ± 2 %, *n* = 6; A2B HOM: 104 ± 4%, *n* = 5; *p* = 0.0008, unpaired *t*-test).

Based on these findings, we wanted to determine whether NMDARs can alter mRNA translation in the absence of ion flux. Hippocampal slices were incubated in ACSF containing actinomycin D ± MK-801 and then labelled with ^35^S methionine/cysteine ± MK-801. During this time slices were briefly stimulated with NMDA or vehicle ± MK-801. We observed that NMDA causes a significant reduction in ^35^S incorporation in the presence of MK-801 to block ion flux during brief stimulation with NMDA (**Fig. 4D**). These findings suggest that NMDARs can alter translational rates in the absence of ion permeation to support spine shrinkage at CA1 synapses. Although it may seem paradoxical that bulk protein synthesis is decreased by stimulating the mTORC1 pathway through GluN2B, these findings are consistent with previous studies in the *Tsc2^+/−^* mouse^35^ and reflect a change in the balance of between pools of mRNA competing for access to ribosomes^36^.

Encouraged by these findings, we next wanted to examine whether switching the GluN2A and GluN2B CTD could influence protein synthesis rates under basal conditions in the absence of exogenous stimulation. We observed that deleting the GluN2B CTD and replacing it with GluN2A (GluN2B^2ACTD^/B2A mice) led to a significant increase in protein synthesis relative to WT slices (**Fig. 4E**). Conversely, eliminating the GluN2A CTD and replacing it with 2B, significantly reduced protein synthesis in GluN2A^2BCTD^ heterozygous and homozygous mice in a gene dose-dependent manner (**Fig. 4F**). Together these findings suggest that the GluN2 CTD regulates basal protein synthesis levels in the hippocampus.

Alterations in global protein synthesis represent a shift in the translating pool of mRNA transcripts^37,38^. Consequently, we expect that manipulating the GluN2 CTDs will impact protein synthesis-dependent phenotypes including synaptic plasticity, circuit excitability and behavior. This has been demonstrated numerous times when studying mutations in core synaptic proteins, such as FMRP, SynGAP and Tuberin^35,38–40^. One of the predominant forms of protein synthesis-dependent synaptic plasticity is mGluR-LTD^41^. Increases in bulk protein synthesis, observed in mouse models of fragile X and *Syngap1* haploinsufficiency, coincide with an elevation in mGluR-LTD that is no longer gated by new protein synthesis^20,40,42^. In contrast, decreases in bulk protein synthesis, as observed in *Tsc2* heterozygous mice, lead to an impairment in mGluR-LTD^35^. Together they define an axis of synaptic pathophysiology, where deviations in protein synthesis can be used to predict disease related cellular abnormalities. Therefore, we hypothesized that protein synthesis-dependent phenotypes may be disrupted in GluN2B^2ACTD^ and GluN2A^2BCTD^ mice, mimicking *Fmr1* KO and *Tsc*2 heterozygous mice, respectively.

To test this hypothesis, we first examined mGluR-LTD in GluN2A^2BCTD^ and GluN2B^2ACTD^ mice. Acute hippocampal slices were prepared from GluN2A^2BCTD^ and GluN2B^2ACTD^ mice and recovered for 3-4 hours before extracellular recordings were performed in CA1. Once a steady baseline was obtained, mGluR-LTD was induced by the acute application of the group 1 mGluR_1/5_ agonist dihydroxyphenylglycine (DHPG; 50 µM, 5 mins^43^), and the slope of fEPSPs were monitored for a further 55 minutes. We observed that DHPG caused a long-lasting depression of fEPSP responses in WT slices, which was exaggerated in the hippocampus of GluN2B^2ACTD^ mice (**Fig. 4G**). In contrast, the magnitude of mGluR-LTD was significantly reduced in GluN2A^2BCTD^ mice (**Fig. 4H**). Thus, GluN2B^2ACTD^ and GluN2A^2BCTD^ show mirror opposite alterations in mGluR-LTD relative to WT littermates, reminiscent of findings observed in mouse models of FXS and TSC, respectively. The finding that the maximal transient depression immediately following application of DHPG did not differ between GluN2 CTD genotypes suggests that these findings are unlikely to be accounted for by differences in mGluR_5_ expression^44^.

### Loss of mNMDAR signaling promotes epileptogenesis in the hippocampus

Previous studies have shown that elevation of basal protein synthesis can lead to epileptogenesis within area CA3 of the hippocampus^45^, which can be mimicked by stimulating mGluR_1/5_ with DHPG. To examine whether the GluN2 CTD modulates excitability, we placed a recording electrode in the pyramidal layer of CA3 in transverse hippocampal slices from GluN2A^(2BCTD)^, GluN2B^(2ACTD)^, and WT littermates. Slices were perfused with ACSF for 10 minutes to obtain a baseline before we monitored spontaneous and epileptiform activity in response to the GABA_A_ antagonist bicuculline (50 µM) or DHPG (50 µM), respectively. In control slices, addition of bicuculline led to short, rhythmic, ictal-like events of <1.5 seconds (**Fig. 5B_i_**). However, in the presence of DHPG ictal-like events progressed to prolonged synchronized discharges (**Fig. 5B_ii_**). Intriguingly, in slices from GluN2B^(2ACTD)^ mice, GABA_A_ blockade alone led to a dramatic increase in burst duration and frequency (**Fig. 5C_i_**), similar to the effect of DHPG (**Fig. 5C_ii_**).

**Figure 5:**
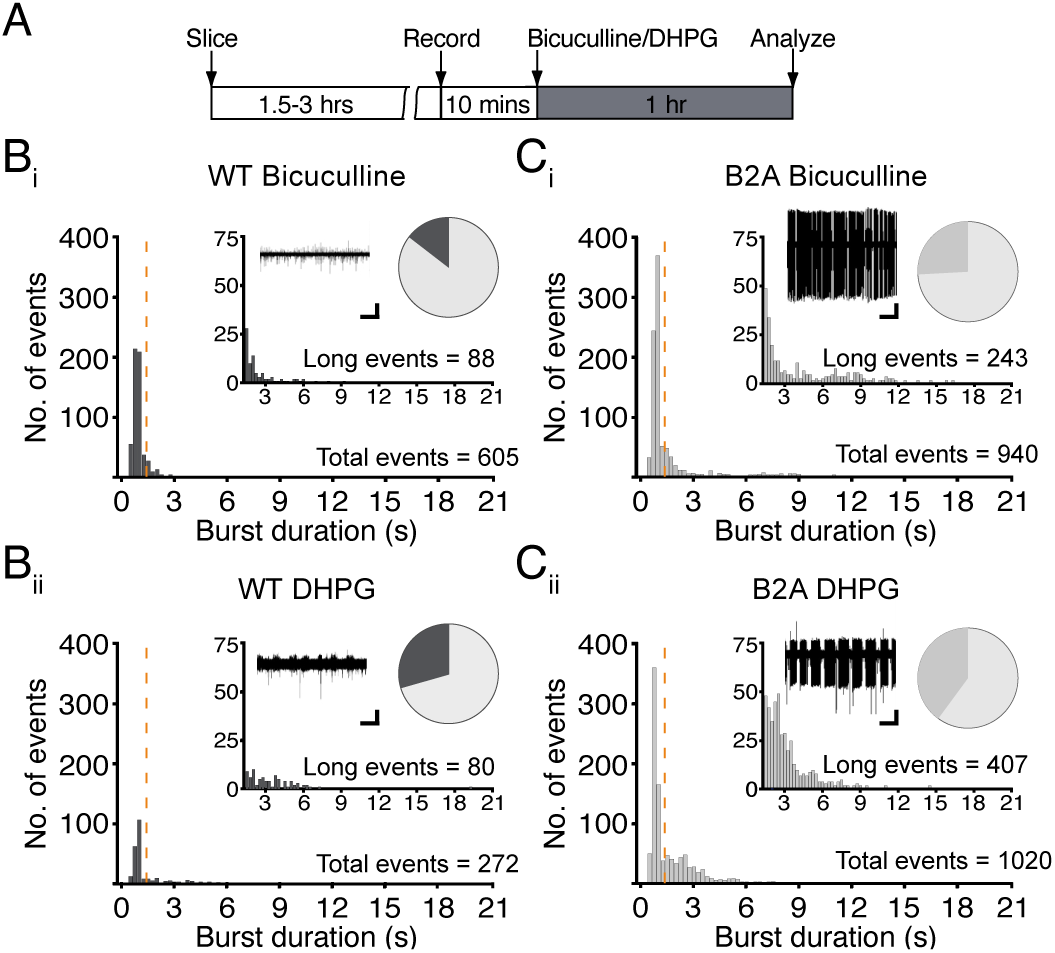
Eliminating mNMDAR signaling by replacing the GluN2B CTD with GluN2A triggers CA3 epileptogenesis. (**A**) Experimental timeline for extracellular recordings in hippocampal CA3. Transverse hippocampal slices were recovered before a recording electrode was placed in the pyramidal layer of CA3. Slices were perfused with ACSF for 10 mins before being treated with either the GABA_A_ receptor antagonist bicuculline (Bic, 50 μM) or the group 1 mGluR_1/5_ agonist DHPG (50 μM) for 1hr. Example traces of extracellular recordings in the presence of bicuculline or DHPG from 50 mins post drug application. (**B_i_**) Extracellular recordings in CA3 pyramidal cell layer of WT hippocampal slices perfused with ACSF containing bicuculline primarily generates short rhythmic events (short events ≤1.5s = 517, long events >1.5s = 88, average burst duration of 605 events = 1.23 ± 0.04s, *n* = 8). (**B_ii_**) Bath application of DHPG generates significantly longer burst events (short events ≤1.5s = 192, long events >1.5s = 80, average burst duration of 272 events = 1.76 ± 0.10s, *n* = 11, *p* < 0.0001 ANOVA). (**C_i_**) GABA_A_ blockade with bicuculline triggers a greater number of epileptiform discharges that are longer in burst duration in CA3 region of hippocampal slices from GluN2B^2ACTD^ (B2A) mice compared to WT mice (B2A: short events <1.5s = 697; long events >1.5s = 243, average burst duration of 940 events = 1.88 ± 0.07s, *n* = 7, *p* < 0.0001, ANOVA). (**C_ii_**) Similarly, burst frequency and duration are increased in B2A slices perfused with DHPG compared to WT mice (B2A: short events <1.5s = 613; long events >1.5s = 407, average burst duration of 1020 events = 1.83 ± 0.0.5s, *n* = 7*, p* < 0.0001, ANOVA). All Scale bars 30s, 50 μV.

In GluN2A^(2BCTD)^ slices, bicuculline had a similar effect to that seen in WT slices (**Fig. 6A-B_i_**), whilst DHPG failed to induce short or long duration events (**Fig. 6A-B_ii_**). Therefore, replacing the GluN2A with the GluN2B CTD blocks mGluR_1/5_ induced epileptogenesis. Together these data suggest that switching the GluN2 CTD leads to a hypo- and hyper-sensitive response to synaptic and mGluR_1/5_ activation in GluN2A^(2BCTD)^ and GluN2B^(2ACTD)^ slices, respectively.

**Figure 6:**
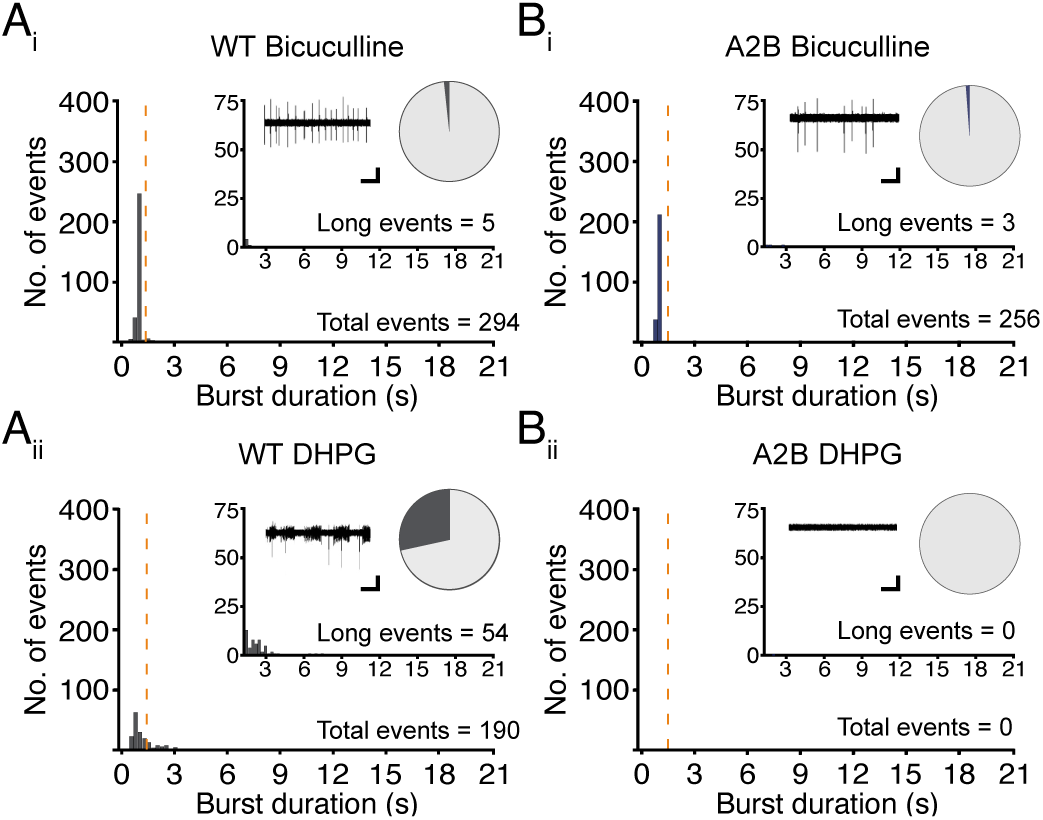
Enhancing mNMDAR signaling by replacing the GluN2A CTD with GluN2B suppresses CA3 epileptogenesis. (**A_i_**) Extracellular recordings in CA3 pyramidal cell layer of WT hippocampal slices perfused with ACSF containing bicuculline primarily generates short rhythmic events (bicuculline: short events <1.5s = 289, long events >1.5s = 5, average burst duration of 294 events = 1.04 ± 0.01s, *n* = 9). (**A_ii_**) Bath application of DHPG generates short rhythmic events that lead to an increase in prolonged discharges (DHPG: short events <1.5s =136, long events >1.5s = 54, average burst duration of 190 events = 1.42 ± 0.07s, *n* = 10) (**A_i_**) Bicuculline induces comparable short and long events in *GluN2A^2BCTD^* (A2B) mice compared to WT littermates (A2B: short events <1.5s = 253; long events >1.5s = 3, average burst duration of 256 events = 1.03 ± 0.01s, *n* = 8). (**B_ii_**) DHPG can no longer induce short or prolonged discharges in A2B hippocampal slices (A2B: short events <1.5s = 0; long events >1.5s = 0, *n* = 7). All Scale bars 30s, 50 μV.

### Switching the GluN2A C-terminal domain with GluN2B normalizes protein synthesis and mGluR-LTD in the *Fmr1^−/y^* mouse

Based on these findings, it appears that GluN2B CTD expression strongly modulates functions related to mGluR_5_-dependent protein synthesis (**Fig. 7A**). Deleting the GluN2B CTD in GluN2B^(2ACTD)^ mice exacerbates phenotypes downstream of mGluR_5_ activation, while genetically enhancing the expression of GluN2B CTD suppresses mGluR_5_-dependent phenotypes in the hippocampus. As there are numerous reports that mGluR_5_ contributes to disease phenotypes in the mouse models of intellectual disability^20,39,46^, we wanted to explore whether manipulating mNMDAR signaling in the *Fmr1* KO mouse could correct some of the core phenotypes in FXS. We introduced the GluN2A^(2BCTD)^ mutation into *Fmr1* KO mice by crossing *Fmr1* heterozygous females with GluN2A^(2BCTD)^ heterozygous males to enhance mNMDAR signaling and measured basal protein synthesis in the resulting progeny (**Fig. 7B**). Our results reveal that the increase in protein synthesis in *Fmr1* KO mice relative to WT littermates is significantly reduced by overexpression of the GluN2B CTD and elimination of the GluN2A CTD with the cross to the GluN2A^(2BCTD)^ mice (**Fig. 7C**).

**Figure 7:**
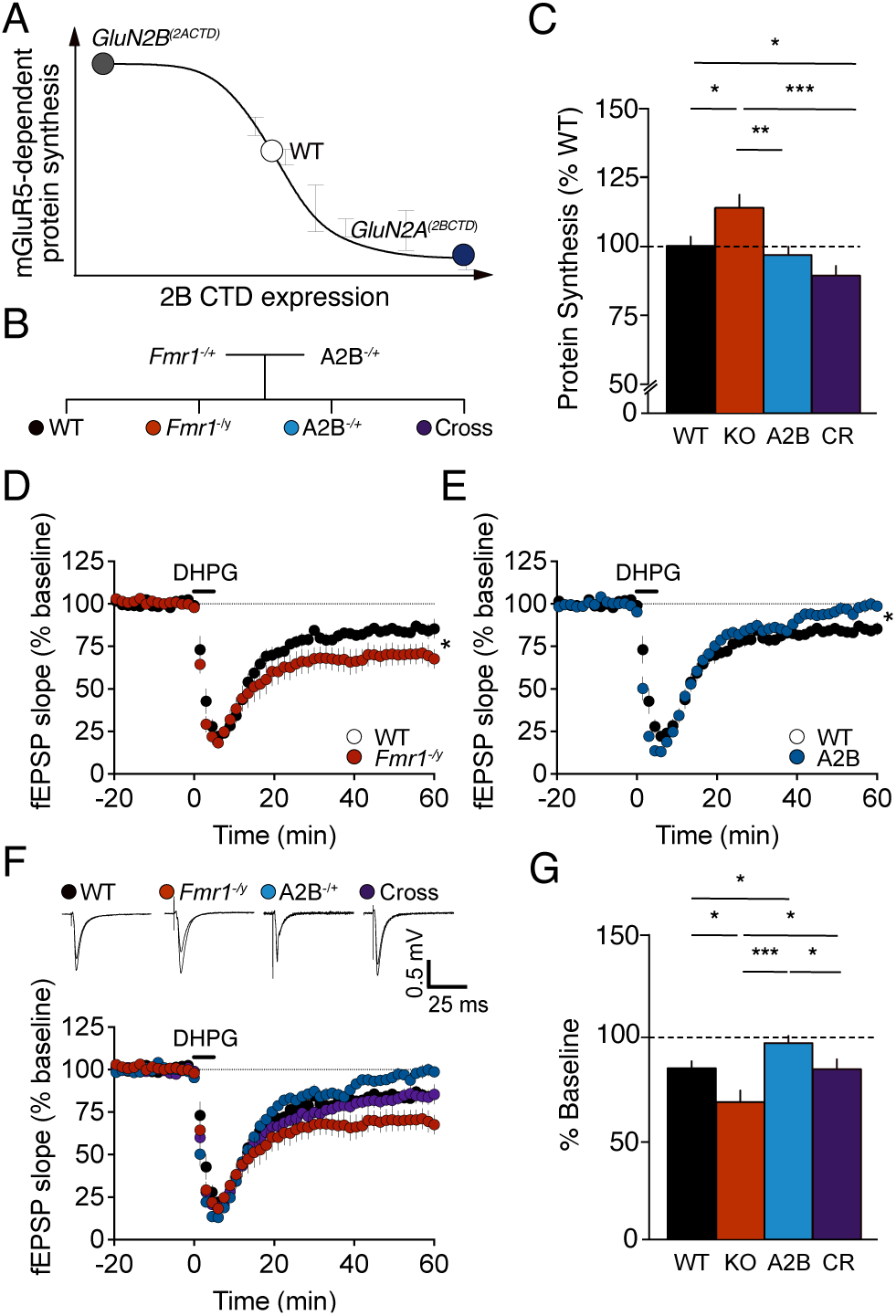
Increasing the expression of GluN2B C-terminal domain normalizes protein synthesis and mGluR-LTD in the *Fmr1^−/y^* mouse. (**A**) Schematic depicting the inverse relationship between GluN2B CTD expression and mGluR_5_ function. (**B**) Genetic rescue strategy in which *Fmr1* heterozygous females are crossed with male *GluN2A^(2BCTD)^* (A2B) heterozygous mice to produce male offspring of four genotypes: WT, *Fmr1* KO, A2B het, and *Fmr1* KO/A2B het cross. (**C**) Replacing the C-terminal domain of GluN2A with GluN2B in *Fmr1* KO reduces elevated protein synthesis rates in the hippocampus (WT: 100 ± 3%, *n* = 21; *Fmr1* KO: 114 ± 5%, *n* = 11; A2B HET: 97 ± 3%, *n* = 13 Cross: 89 ± 3%, *n* = 14; ANOVA A2B genotype *p* = 0.0002). (**D**) The magnitude of mGluR-LTD in hippocampal CA1 is exaggerated in *Fmr1* KO mice relative to WT littermates (WT: 85 ± 3%, *n* = 10; *Fmr1* KO: 70 ± 6%, *n* = 10; unpaired *t*-test, *p* = 0.029). (**E**) in contrast, mGluR-LTD is significantly impaired in *GluN2A^(2BCTD)^* heterozygote mice (A2B: 98 ± 3%, n = 8, unpaired *t*-test, *p* = 0.013). (**F**) Introduction of the *GluN2A^(2BCTD)^* mutation in *Fmr1* KO mice restores elevated mGluR-LTD to WT levels (Cross: 85 ± 4%, *n* = 9, unpaired *t*-test, *p* = 0.99). (**G**) Comparison of all four genotypes reveals a suppressive effect of the *GluN2A^(2BCTD)^* heterozygous mutation, lowering mGluR-LTD in the *Fmr1* KO hippocampus to WT levels (ANOVA A2B/Fmr1 genotype *p* = 0.003).

The correction of protein synthesis with the introduction of the GluN2A^2BCTD^ knockin mutation led us to speculate that other pathological phenotypes could be restored in *Fmr1* KO mice. To examine this, we measured mGluR-LTD in the *Fmr1* KO x GluN2A^(2BCTD)^ cross. Consistent with previous studies, the acute application of DHPG (50 µM, 5 mins) revealed an exaggerated mGluR-LTD in *Fmr1* KO mice relative to WT littermates (**Fig. 7D**). The GluN2A^(2BCTD)^ heterozygous mutation led to a small but significant suppression of mGluR-LTD when compared to WT mice (**Fig. 7E**). Importantly, the *Fmr1* KO x GluN2A^2BCTD^ cross exhibits a magnitude of LTD indistinguishable from WT mice (**Fig. 7F**). Together, our results show that the enhancement of GluN2B CTD expression through the introduction of the GluN2A^2BCTD^ mutation is sufficient to correct excessive protein synthesis and exaggerated mGluR-LTD in the *Fmr1* KO hippocampus.

### Dysregulated protein synthesis and audiogenic seizures in *Fmr1^−/y^* mice are corrected by modulating GluN2B

Previous studies in *Fmr1* KO mice have shown that correction of altered protein synthesis is highly predictive of improvements in structural, circuit, and behavioral phenotypes^36,39,46,47^. According to our model, augmenting GluN2B signaling should ameliorate FXS phenotypes. Glyx-13 has recently been shown to preferentially target GluN2B containing receptors at an allosteric site distinct from the glutamate and glycine binding sites^48,49^. Furthermore, it is known to be a cognitive enhancer in several learning and memory paradigms^50–52^. To test the potential utility of augmenting NMDAR function in fragile X, we treated *Fmr1* KO and WT hippocampal slices with Glyx-13 (0.1 μM) and measured protein synthesis levels (**Fig. 8A**). Once again, we observed a significant increase in ^35^S incorporation in *Fmr1* KO mice relative to WT littermates under basal conditions. Importantly, pretreatment with Glyx-13 normalized excessive protein synthesis in *Fmr1* KO mice to WT levels under these conditions (**Fig. 8B**). This result supports the hypothesis that positively modulating GluN2B under tonic conditions restores one of the core deficits in *Fmr1* KO hippocampus.

**Figure 8:**
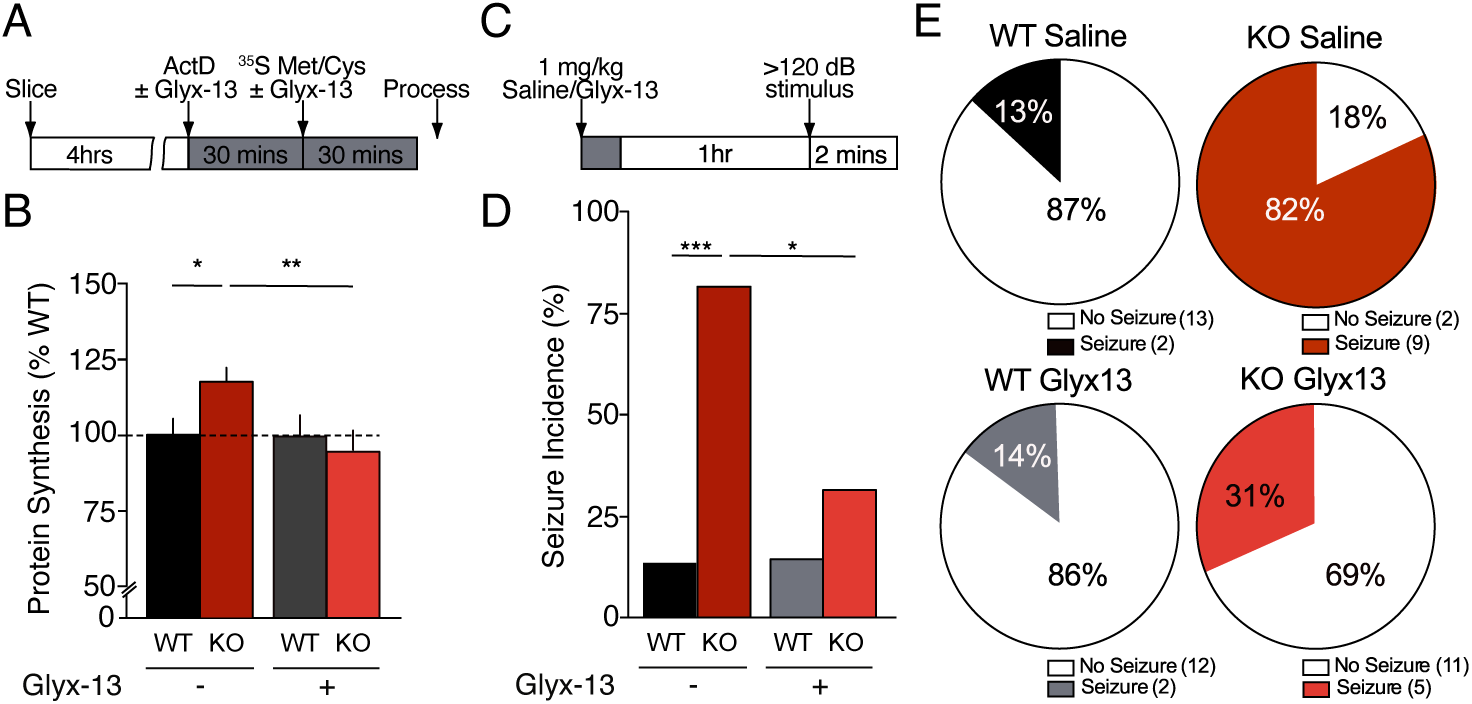
Dysregulated protein synthesis and audiogenic seizures in *Fmr1^−/y^* mice are corrected by modulation of mNMDAR signaling. (**A**) Experimental timeline for metabolic labeling in the presence of the NMDA partial agonist Glyx-13. (**B**) Glyx-13 (0.1 μM) restored elevated protein synthesis in *Fmr1* KO mice to WT levels (WT vehicle: 100 ± 6%; KO vehicle: 118 ± 5%; WT Glyx-13: 99 ± 7%; KO Glyx-13: 94 ± 5%; ANOVA *p* = 0.034, *n* = 8). (**C**) Experimental timeline for the *in vivo* treatment of WT and *Fmr1* KO mice with Glyx-13 (1 mg/kg) prior to audiogenic seizure (AGS) testing. Mice were injected with either saline or Glyx-13 (1 mg/kg) 1 hr prior to AGS testing. Seizures were scored for wild running, clonic seizure, tonic seizure and death. (**D, E**) AGS incidence is significantly increased in saline treated *Fmr1* KO mice relative to WT littermates (WT saline: 13%, *n* = 15; KO saline: 82%, *n* = 11; Fisher’s exact test *p* = 0.0009). Treatment with Glyx-13 had no effect on the incidence of seizures in WT mice, but significantly reduced AGS incidence in *Fmr1* KO mice (WT Glyx-13: 14%, n = 14; KO Glyx-13: 31%, *n* = 16, Fisher’s exact test *p* = 0.018).

In addition to an increase in spine density and hyperexcitability within the hippocampal circuit, another consequence of altered protein synthesis in *Fmr1* KO mice is enhanced susceptibility to audiogenic seizures (AGS)^39,46,47^. Encouraged by our initial findings we next wanted to determine whether Glyx-13 could reduce the incidence of AGS in *Fmr1* KO mice *in vivo. Fmr1* KO and WT littermates (P19-25) mice were injected with either saline or Glyx-13 (1mg/kg). After 1 hr, mice were transferred to a chamber and habituated for 1 minute before they were exposed to a 120dB stimulus for 2 minutes (**Fig. 8C**). As expected, saline-treated *Fmr1* KO mice exhibited an increased susceptibility to seizures relative to WT littermates (**Fig. 8D, E**). However, a single dose of Glyx-13 (1mg/kg) was able to significantly reduce seizure susceptibility, without impacting the incidence of seizures in WT mice. These findings further support the hypothesis that positively modulating GluN2B signaling can rescue both i*n vitro* and *in vivo* phenotypes in *Fmr1* KO mice.

## DISCUSSION

The current study reveals that GluN2B is necessary and sufficient for mNMDAR signaling in the absence of ion flux, and that this signaling can mitigate core fragile X phenotypes linked to altered protein synthesis regulation. In addition, the data clarify the roles of GluN2A and GluN2B subunits in initiating NMDAR-dependent LTD and structural plasticity in CA1.

### NMDAR molecular motifs involved in non-ionotropic structural and ionotropic functional plasticity in CA1

We have previously shown that changes in the length and volume of dendritic spines in response to NMDA proceeds independently of ion flux and is dependent on mTORC1 and *de novo* protein synthesis^16^. Conversely, LTD of synaptic transmission initiated by NMDA occurs in the absence of mTORC1 activity or new protein synthesis but requires ion flux through the receptor channel. In the current study we find that structural and functional plasticity in CA1 can be further dissociated by their dependence on GluN2 subunits. LTD, as measured by a change in synaptic transmission evoked by stimulation of the Shaffer collaterals, is dependent on both the GluN2A and GluN2B subtypes, while non-ionotropic signaling as reported by spine shrinkage is solely dependent on receptors containing GluN2B.

We found that pharmacologically limiting ion flux through GluN2A-containing assemblies, or deleting the *Grin2A* gene (constitutively or conditionally), reduces the magnitude of NMDAR-LTD. However, LTD was fully abolished only when the LBD of GluN2B was blocked. Combined, these data suggest that both the GluN2B/GluN1 di-heteromeric and GluN2A/GluN2B/GluN1 tri-heteromeric assemblies contribute to LTD at CA1 synapses, whereas GluN2A/GluN1 di-heteromeric receptors do not. The GluN2B subunit is also crucial for non-ionotropic signaling reported as spine shrinkage. Structural plasticity was eliminated by antagonizing the LBD of GluN2B, genetically deleting this receptor subunit, or swapping out its CTD. Indeed, because deletion of GluN2A has no quantitative effect on spine shrinkage in response to NMDA, the data suggest that the GluN2B di-heteromeric receptors can account for the full effect of mNMDAR signaling.

Previous work has implicated CAMKII, p38 MAPK and neuronal nitric oxide synthase signaling in driving actin depolymerisation and the retraction of dendritic spines^6,8^ as well as gating LTP-associated spine growth associated with long-term potentiation^9^. Our results indicate that GluN2B di-heteromeric assemblies are sufficient to support spine shrinkage caused by mNMDAR activation, and that the GluN2B LBD and CTD are critical structural motifs. Considered with the previous study by Thomazeau et al (2020), who also used NMDA to trigger spine shrinkage, the data indicate that GluN2B signals through mTORC1 to modulate protein synthesis that serves as a gate for structural plasticity.

How might the GluN2B CTD mediate its non-ionotropic functions and couple to downstream pathways controlling spine morphology and translation? Biochemical and genetic studies have demonstrated that the GluN2B CTD plays an essential role in the supramolecular assembly of NMDARs with scaffold proteins and signaling enzymes, which are key regulators of structural and functional plasticity^53^. GluN2B di- and tri-heteromers assemble into 1.5MDa supercomplexes whereas GluN2A di-heteromers can only assemble into 800kDa complexes lacking MAGUKs and signaling molecules. Thus, the physiological functions observed for the GluN2B CTD could arise from its role in positioning of the receptor with its effector proteins and/or a direct metabotropic signaling function exerted on the signaling complex.

### Regulation of protein synthesis by mNMDAR signaling

Further examination of mNMDAR signaling reveals that the GluN2B subtype can bidirectionally regulate protein synthesis. Brief NMDA application reduces bulk protein synthesis while antagonizing the NMDAR LBD under basal conditions causes a significant elevation. Importantly, modulation of translational rates occurs in the presence of the channel blocker MK-801, suggesting that non-ionotropic mNMDAR signaling represents an important pathway by which synaptic activity regulates protein synthesis.

Consistent with this conclusion, we find that augmentation of mNMDAR signaling by replacing the GluN2A CTD with the GluN2B CTD mimics the effect of agonist application on protein synthesis in WT hippocampus. Conversely, elimination of mNMDAR signaling by swapping out the GluN2B CTD mimics the effect of D-AP5 on WT protein synthesis. The functional significance of protein synthesis modulation by mNMDAR signaling through GluN2B is suggested by the impact on LTD induced by activating mGluR_5_ with DHPG^43^. DHPG stimulates new protein synthesis that is required for induction of mGluR-LTD in WT mice^39,41,43^. DHPG-induced mGluR-LTD is augmented in slices from the GluN2B^(2ACTD)^ mice with increased protein synthesis rates, and it is reduced in slices from the GluN2A^(2BCTD)^ mice with reduced basal protein synthesis. Going forward, it will be of great interest to investigate how the well-known effects of age and sensory experience on NMDAR subunit composition impact protein synthesis regulation and its functional consequences during development^54^.

Our findings of reduced global protein synthesis in response to mNMDAR activation are consistent with earlier reports in cortical cultures which demonstrated that the GluN2B subunit suppresses protein synthesis^55,56^. Interestingly, global protein synthesis is also reduced in a mouse model of tuberous sclerosis in which mTORC1 signaling is hyperactive^35^. In this case, suppression of overall protein synthesis reflects a decrease in translation of terminal oligopyrimidine (TOP) containing mRNA transcripts and an increase in FMRP-binding targets that tend to encode large synaptic proteins^38^. When considered with the finding that mNMDAR-induced structural plasticity is blocked by the mRNA translation inhibitor cycloheximide, these results suggest that GluN2B triggers a shift in the translation pool rather than stalling mRNA translation *per se*. Based on these findings, we propose that agonist binding to the GluN2B LBD triggers a conformational change in the CTD of GluN2B di-heteromers to activate mTORC1 signaling and initiate mRNA translation and the synthesis of proteins that support the depolymerisation of the actin cytoskeleton and the retraction of dendritic spines. It will be of great interest to identify actively translating mRNAs downstream of non-ionotropic mNMDAR signaling and understand how they contribute to structural plasticity at CA1 synapses.

### Mimicry and correction of core cellular phenotypes in *Fmr1^−/y^* mice

Recently, it has been reported that elevated protein synthesis in *Fmr1* KO mice reflects a length-dependent shift in the translating pool of mRNAs^37^. There is an over-translation of short mRNA transcripts encoding ribosomal proteins at the expense of longer mRNAs encoding synaptic proteins and FMRP targets. Consequently, synaptic and circuit paradigms that are normally dependent on *de novo* protein synthesis are disrupted in the *Fmr1* KO and persist in the presence of protein synthesis inhibitors^16,42,57^. We previously found that this also applies to NMDA-induced spine shrinkage in *Fmr1* KO mice, which unlike WT, proceeds in the presence of cycloheximide^18^.

Here we show that replacing the GluN2B CTD with GluN2A, and thus eliminating mNMDAR signaling, leads to increased protein synthesis and spine density, exaggerated mGluR-LTD in CA1, and epileptiform activity in CA3. These differences in hippocampal structure and function phenocopy observations in the *Fmr1^−/y^* mouse^20,47,58^. In contrast, overexpression of the GluN2B CTD reduces both LTD and the increase in neuronal excitability triggered by mGluR_5_ activation.

Collectively, these data demonstrate that non-ionotropic mNMDAR signaling strongly modulates the intracellular consequences of activating mGluR_5_, which have been linked to pathophysiology in several neurodevelopmental disorders that, in addition to fragile X, include tuberous sclerosis complex^35,59^, Rett syndrome^60^, chromosome 16p11.2 microdeletion syndrome^61^, and autism^62^.

Numerous prior studies have shown that selectively inhibiting mGluR_5_ or its downstream signaling normalizes protein synthesis and rescues a multitude of synaptic and behavioral phenotypes in the *Fmr1* KO^63^. Similarly, enhancing mTORC1 signaling through the introduction of the *Tsc2* mutation corrects synaptic pathophysiology in FXS^35^. This motivated us to try and reverse the core pathophysiology associated with *Fmr1* KO mice, by increasingly mNMDAR signaling through the introduction of *GluN2A^2BCTD^* mutation. In the absence of FMRP, increasing the expression of the GluN2B CTD reduced exaggerated protein synthesis, mGluR-LTD, and epileptiform activity in the hippocampus of *Fmr1* KO mice.

Allosteric augmentation of mNMDARs, in principle, could improve synaptic regulation of mTORC1 and by doing so restore the normal balance of high- and low-efficiency mRNA translation in fragile X^64,65^. In a preliminary test of this hypothesis, we observed that Glyx-13, a positive modulator of NMDARs^51,66^, corrected the protein synthesis phenotype in *Fmr1^−/y^* hippocampus. Encouraged by these findings, we also investigated the consequence of modulating NMDARs on audiogenic seizures, an *in vivo* phenotype in *Fmr1^−/^*^y^ mice that is associated with altered protein synthesis regulation^67^. A single dose of Glyx-13^68^ was sufficient to significantly reduce the incidence of seizures. Glyx-13 acts on NMDARs at an allosteric site distinct from the glutamate and glycine binding sites^48^. It remains to be determined if the effect of this compound on AGS is independent of effects on NMDAR ion flux.

Mutations in GluN2B have been linked to numerous types of epilepsy including epileptic encephalopathies, focal epilepsy, partial seizures and infantile spasms^69–71^. In addition, deletion of GluN2B leads to an increase in the number of excitatory inputs on to pyramidal neurons in the prefrontal cortex^56^. In the current study we also detected an increase in CA1 spine density in the absence of the GluN2B CTD and, in CA3, we find prolongation of epileptiform bursts in response to mGluR_5_ activation. Together, these findings demonstrate that the GluN2B subunit makes a critical contribution to circuit refinement, protein synthesis regulation, and circuit excitability in the juvenile hippocampus. Identifying compounds that can selectively modulate this unique mode of signaling may prove to be a highly valuable target in treating neurodevelopmental disorders.

## Author contributions

SAB, AT, EKO and MB designed research; SAB, AT, PSBF, AJH, and MJH performed research; SAB, AT, and MJH analyzed data; SAB and MFB wrote the paper; MFB and EKO supervised the project. The authors declare no conflict of interest.

## Acknowledgements

Special thank you to Joseph Moskal for the helpful advice on Glyx-13. The authors are grateful to the FRAXA foundation and The Picower Institute Innovation Fund for supporting this research. Additional support was provided by NIH grant R21NS123499.

